# *Acinetobacter baumannii* and cefiderocol between *cidality* and adaptability

**DOI:** 10.1101/2022.05.05.490855

**Authors:** Stefano Stracquadanio, Carmelo Bonomo, Andrea Marino, Dafne Bongiorno, Grete Francesca Privitera, Dalida Angela Bivona, Alessia Mirabile, Paolo Giuseppe Bonacci, Stefania Stefani

## Abstract

Among the bacterial species included in the ESKAPE group, *Acinetobacter baumannii* is of great interest due to its intrinsic and acquired resistance to many antibiotic classes and its ability to infect different body districts. Cefiderocol is a novel cephalosporin active against Gram-negative bacteria with promising efficacy on *A. baumannii* infections, but some studies have reported therapeutic failures even in the presence of susceptible *A. baumannii* strains.

This study aims to investigate the interactions between cefiderocol and ten *A. baumannii* strains with different susceptibility profiles to this drug. We confirmed diverse susceptibility profiles in the strains, with resistance values close to the EUCAST-proposed breakpoints. MBC/MIC (minimal bactericidal concentration/minimal inhibitory concentration) ratios, demonstrated bactericidal activity of the drug; on the other hand, bacterial regrowth was evident after exposition to cefiderocol, as were changes in the shape of colonies and bacterial cells. A switch to a non-susceptible phenotype in the presence of high cefiderocol concentrations was found as adaptation mechanisms implemented by these *A. baumannii* strains to overcome the *cidal* activity of this antibiotic, also confirmed by the presence of heteroresistant, unstable subpopulations. As our isolates harbored numerous β-lactamase genes, β-lactamase inhibitors showed the ability to restore the antimicrobial activity of cefiderocol regardless of the different non-susceptibility levels of the tested strains.

These in vitro results, can sustain the concept of using combination therapy to eliminate drug-adapted subpopulations and regain full cefiderocol activity in this difficult-to-treat species.

## INTRODUCTION

*Acinetobacter baumannii* is a major clinical threat, both because of its established role as an important pathogen associated with nosocomial infections at various body sites and the challenge posed by the very limited treatment options available, due to its high intrinsic and acquired antimicrobial resistance. The latter entails, in many cases and for different antibiotics, characteristics of instability, related - as recently demonstrated – to frequent genomic rearrangements involving gene amplification, MGE acquisition, resistome and phage profiling changes [1,2].

The recent introduction of the novel siderophore-conjugated cephalosporin cefiderocol, whose spectrum includes *A. baumannii*, has raised great expectation in therapy [3]. The molecule is composed of a siderophore component that binds iron and uses active iron transport for drug entry into the bacterial periplasmic space. The cephalosporin moiety is the active antimicrobial component, structurally resembling a hybrid between ceftazidime and cefepime. Like other β-lactam agents, the main bactericidal activity of cefiderocol occurs through the inhibition of bacterial cell wall synthesis via binding of penicillin-binding proteins (PBPs) and inhibition of peptidoglycan synthesis, leading to cell death [4].

Several clinical studies have pointed out that this drug had similar efficacy to the best therapies available for infections caused by Gram-negative bacteria, but a higher rate of all-cause mortality has been described in the subset of *A. baumannii* infections [5], despite a minimal inhibitory concentration (MIC) of 4 mg/L. These authors indicated heteroresistance as a possible responsible for these failures [5,6], especially if bacteria are classified as susceptible by standard antibiotic susceptibility testing (AST) [1].

Heteroresistance is a poorly understood mechanism of survival in the presence of antibiotics and is defined as the presence of subpopulations with MIC values higher (variably defined as equal to or more than two- to eight-fold) than in the main population, i.e., a susceptible bacterial isolate could harbor a minority of resistant subpopulations. Despite the discovery of a high prevalence of this mechanism in many different species, both Gram-positive and Gram-negative, with respect to many antibiotics [1], the majority of these subpopulations were considered unstable. Heteroresistance in *A. baumannii* was already described for different classes of antibiotics, including colistin, aminoglycosides, imipenem, meropenem, and tigecycline [7–10].

Due to the problem of reproducibility of broth microdilution (BMD) testing [11] as well as reported errors in disk diffusion [12], we performed some *cidality* tests and population analysis profiling (PAP) on a subset of isolates representative of the entire sample with the aim to determine the best *in vitro* conditions to test cefiderocol activity on *A. baumannii* strains with different degrees of susceptibility to this drug. Our results demonstrated that this species has a great potential for heteroresistance, despite an initial result of bactericidal activity of cefiderocol obtained from the evaluation of the MBC/MIC (minimal bactericidal concentration/minimal inhibitory concentration) ratio. Furthermore, these subpopulations, which in some cases are the result of antibiotic induction, were not stable at all, as they returned to the initial MIC after two passages in an antibiotic-free medium.

Interestingly, the addition of avibactam and sulbactam eliminates all subpopulations.

## MATERIAL AND METHODS

### Bacterial strains and culture conditions

Three laboratory-adapted and seven clinical MDR *A. baumannii* strains were selected for this study. Laboratory-adapted *A. baumannii* ATCC 19606 and ATCC 17978 were commercially obtained from the American Type Culture Collection (Manassas, VA) and *A. baumannii* ACICU was provided courtesy of Prof. Paolo Visca (Roma Tre University, Italy), while clinically significant strains were recently isolated from bronchial aspirates of cefiderocol (FDC)-naïve hospitalized patients, and FDC testing was requested by the infectious disease specialist. The clinical isolates were identified as *A. baumannii* by Matrix-Assisted Laser Desorption Ionization-Time of Flight (MALDI- TOF) mass spectrometry (Bruker Daltonics, Billerica, MA, USA) [13]. The strains were grown in

McConkey agar plates (Oxoid, UK) at 37 °C for 18 h and stored in Trypticase Soy Broth (TSB) (Oxoid, UK) plus 15% glycerol at −80 °C until further analysis.

### Antibiotic susceptibility testing

Meropenem (MEM), colistin (COL), and FDC *in vitro* susceptibility was evaluated in accordance with the EUCAST guidelines [14,15].

MEM trihydrate and COL sulfate salt powders (Merck/Sigma-Aldrich, Germany) were used to assess MICs by BMD in cation-adjusted Mueller Hinton II broth (CAMHB) (Becton, Dickinson and Company, MD, USA).

FDC disks containing 30 μg of antibiotic (Liofilchem, Italy) and FDC powder (SHIONOGI, Japan) were provided courtesy of SHIONOGI. Disk diffusion (DD) assays were performed on Mueller Hinton agar plates (MHA) (Oxoid, UK), while BMD MIC determination was performed on both CAMHB and iron-depleted cation-adjusted Mueller Hinton II broth (ID-CAMHB), the latter as indicated in the EUCAST guidelines, prepared using the Chelex® 100 resin (Bio-Rad Laboratories, CA, USA), as reported by Hackel et al. [11]

All experiments were performed in triplicate and the geometric mean was then calculated.

Due to the lack of actual breakpoints for FDC susceptibility in *A. baumannii*, strains with an inhibition zone with a diameter ≥17 mm were considered susceptible, whereas a diameter <17 mm was indicative of non-susceptible *A. baumannii* strains. Similarly, strains with a MIC value >2 were considered non-susceptible to FDC [14]. The minimal bactericidal concentration (MBC) was determined by plating the dilution representing the MIC as well as two less concentrated and five more concentrated dilutions from the same MIC assay on MHA plates and counting viable colonies. MBC was defined as the lowest concentration that demonstrated a reduction of 99.9% in CFU/ml compared to the starting inoculum. The MBC/MIC ratio was calculated from MIC assays performed in ID-CAMHB to see whether FDC had bactericidal or bacteriostatic activity on *A. baumannii* strains; a ratio of ≤4 was indicative of a bactericidal effect [16].

### Population analysis profile (PAP)

Population analysis profile was performed and evaluated as described previously by Choby et al. and Band et al. [6,17] with minor modifications. Briefly, a well isolated colony from a McConkey agar plate was inoculated in 3 mL of tryptic soy broth (TSB) (Oxoid, UK) and incubated overnight at 37 °C with shacking. The bacterial culture was serially diluted in sterile H_2_O from 10^-1^ to 10^-6^, and 50 μL of each dilution was plated in duplicate on MHA plates containing 0 (free), 1, 2, 4, 8, 16, or 32 mg/L of FDC (limit of quantification 20 CFU/mL). Colonies were enumerated after 24 and 48 hours of growth at 37 °C, and only plates with 10 to 300 colonies were considered for subsequent analysis. Isolates were classified as resistant if the number of colonies that grew at the breakpoint concentration (i.e., 2 mg/L) was ≥50% of those growing on antibiotic-free plates. If an isolate was not resistant, it was classified as heteroresistant if the number of colonies that grew at the most concentrated dilution of FDC (i.e., 32 mg/L) was at least 1:10^6^ of those growing on antibiotic-free plates. Isolates that were classified as neither resistant or heteroresistant were classified as susceptible. Student’s t-tests of the ratios between the number of CFU/mL grown on FDC 2 mg/L MHA plates and 50% of the colonies grown on FDC-free plates were performed using GraphPad Prism version 8.0.0 for MacOS (GraphPad Software, California, USA) and only *p-values* ≤0.05 were considered statistically significant. The morphology of the bacteria exposed to different concentration of FDC was evaluated by optical microscopy.

### Cefiderocol non-susceptibility induction

Starting from the PAP assay, one colony was recovered from a MHA plate with 32 mg/L of FDC and streaked on a fresh FDC 32 mg/L MHA plate and inoculated in 3 mL of ID-CAMHB with 32 mg/L of FDC. If the strain was able to grow in the presence of such a high concentration of antibiotic, a disk diffusion test and MIC evaluation were performed as described above to evaluate a potential change in the susceptibility profile of the strain. The same tests were repeated after culturing the previously induced strain on FDC-free MHA plates for 2 days to determine whether the acquired non-susceptibility to the drug was stable. The morphology of the induced bacteria, either exposed or not to the highest concentration of FDC, was evaluated by optical microscopy.

### Evaluation of the synergistic activity of avibactam and sulbactam

The synergistic activity of these two β-lactamase inhibitors (BLIs) in combination with FDC was investigated by disk diffusion test or e-test using disks of ceftazidime (CAZ) and ceftazidime/avibactam (CZA) (Oxoid, UK) - containing 4 μg of avibactam - or strips of ampicillin (AMP) and ampicillin/sulbactam (SAM) (Liofilchem, Italy) - containing 4 mg/L of sulbactam - placed both on FDC-free MHA plates and MHA plates containing 32 mg/L or 1 mg/L of FDC previously inoculated with one FDC-non susceptible induced strain (FDC MIC >32 mg/L) or strains with a FDC MIC >1 mg/L, respectively. *A. baumannii* ATCC 17978 was used as FDC 1 mg/L activity control. The comparison between the inhibition obtained by CAZ and CZA or AMP and SAM in the plates with and without FDC allowed us to evaluate the efficacy of the BLIs.

## RESULTS

The strains included in study are described in Tables 1a and 1b. All isolates were sequenced by NGS (data not shown) and belong to ST 2 (n.8 strains), ST 52 (n.1 strain), ST 77 (n.1 strain). All isolates contained several OXA gene alleles (Table 1a), including the acquired OXA23 (6 out of 7 clinical isolates). All ten strains had AmpC (ADC variants such as ADC2, ADC25, ADC30, ADC73). The analysis of the mutations in the septum formation penicillin binding protein 3 (PBP3 - proposed as the main FDC target), showed only a non-synonymous mutation in Abau2 leading to the A515V amino acid change, and two non-synonymous mutations in ACICU producing A346V and H370Y amino acid variations. Moreover, we found seven (ATCC 19606), ten (all the clinical strains) or 16 (ACICU) synonymous mutations in the same gene (reference genome *Acinetobacter baumannii* ATCC 17978, GenBank code CP000521.1, locus tag A1S_3204).

**Table 1.** Sequence type, β-lactamase genes and antibiotic resistance profile of the study sample.

| a) | | Geometric mean $\sqrt[n]{(x_1 \cdot x_2 \cdot x_3 \dots x_n)}$ | | | | | | | | | | b) | | |
| --- | --- | --- | --- | --- | --- | --- | --- | --- | --- | --- | --- | --- | --- | --- |
| Genomic analysis |  |  |  | MIC mg/L |  | MH Agar |  | CAMHB | ID-CAMHB |  |  | PAP |  |  |
| Strain | ST | *blaOxa<sub>n</sub>* | *blaADC<sub>n</sub>* | MEM | COL | DD<br>mm | S/HR/NS<sup>‡</sup> | MIC<br>mg/L | MIC<br>mg/L | MBC<br>mg/L | MBC/<br>MIC | CFU/mL<br>2mg·L<sup>-1</sup>/Free | CFU/mL<br>32mg·L<sup>-1</sup>/Free | S/HR/R<sup>†</sup> |
| Abau1 | 2 | Oxa<sub>23,66,82,115,174,175,177,201</sub> | Adc<sub>25</sub> | 64 R | 1 S | 12 | NS | 4 | 8 | 16 | 2 | >50% | >1/10<sup>6</sup> | HR |
| Abau2 | 2 | Oxa<sub>23,66</sub> | Adc<sub>73</sub> | 64 R | 2 S | 24 | S | 1 | 0.5 | 2 | 4 | <50% | >1/10<sup>6</sup> | HR |
| Abau3 | 2 | Oxa<sub>23,66,82,115,174,175,177,201</sub> | Adc<sub>25</sub> | 64 R | >64 R | 17 | S | 1 | 4 | 8 | 2 | >50% | >1/10<sup>6</sup> | HR |
| Abau4 | 52 | Oxa<sub>66,72,82,897</sub> | Adc<sub>30</sub> | >64 R | 8 R | 16 | NS | 8 | 1 | 2 | 2 | >50% | >1/10<sup>6</sup> | HR |
| Abau5 | 2 | Oxa<sub>23,66,82,115,174,175,177,201</sub> | Adc<sub>25</sub> | >64 R | 2 S | 15 i.c. | NS/HR | 4 | 4 | 4 | 1 | >50%* | >1/10<sup>6</sup> | R |
| Abau6 | 2 | Oxa<sub>23,66,82,115,174,175,177,201</sub> | Adc<sub>25</sub> | >64 R | 1 S | 15 i.c. | NS/HR | 8 | 4 | 4 | 1 | >50% | >1/10<sup>6</sup> | HR |
| Abau7 | 2 | Oxa<sub>23,66,82,115,174,175,177,201</sub> | Adc<sub>25</sub> | >64 R | 2 S | 15 | NS | 8 | 16 | 32 | 2 | >50% | >1/10<sup>6</sup> | HR |
| ACICU | 2 | Oxa<sub>20,66</sub> | Adc<sub>25</sub> | 64 R | 1 S | 17 | S | 8 | 2 | 8 | 4 | >50% | >1/10<sup>6</sup> | HR |
| ATCC 19606 | 2 | Oxa<sub>98</sub> | Adc<sub>2</sub> | 16 R | 1 S | 26 | S | 0.12 | 0.06 | 0.5 | 8 | <50%* | >1/10<sup>6</sup> | HR |
| ATCC 17978 | 77 | Oxa<sub>95</sub> | Adc<sub>25</sub> | 1 S | 1 S | 22 | S | 0.25 | 0.06 | 0.25 | 4 | <50%* | <1/10<sup>6</sup> | S |

<sup>‡</sup>characterized as susceptible or non-susceptible in accordance with the EUCAST breakpoints for cefiderocol and AST: a zone diameter $\geq 17$ mm is typical for isolates with MIC values $\leq 2$ mg/L
(*Enterobacterales*, *P. aeruginosa*, and PK/PD breakpoint) [14]; the presence of colonies within a zone of clearing indicates a possible heteroresistance phenotype [17]
<sup>†</sup>characterized as susceptible, heteroresistant or resistant following the published protocol [6,16]
\**p*-value $\leq 0.05$ (Student's t-test)
i.c.: colonies within the zone of clearing
S: susceptible; HR: heteroresistant; NS: non-susceptible; R: resistant
MEM: meropenem; COL: colistin

### MIC, MBC and *cidality*

Table 1a shows the MEM and COL MIC values together with the FDC *in vitro* susceptibility profile in terms of MIC, MBC and MBC/MIC ratio to determine drug *cidality*. In accordance with the EUCAST criteria for susceptibility categorization, all strains but *A. baumannii* ATCC 17978 were resistant to MEM, while only two strains (namely Abau3 and Abau4) were COL-resistant.

FDC DD results highlighted that two clinical *A. baumannii* strains and the three controls were susceptible to FDC, while four clinical strains were non-susceptible, with two strains showing colonies within the FDC inhibition zones, which in itself demonstrates the presence of subpopulations, as reported by Sherman et al. [18].

MIC assays were performed in triplicate in both CAMHB and ID-CAMHB, and their geometric means were compared with DD tests to evaluate concordance. In seven strains, inhibition zone and MIC values correlate, whereas two clinical strains (Abau3, Abau4) showed discordance between DD and MIC in ID-CAMHB and the laboratory-adapted ACICU strain displayed different susceptibility phenotypes when tested by DD and in CAMHB for MIC evaluation, respectively. Of note, all discrepancies were observed when the inhibition zone was 17 mm (the susceptibility breakpoint) in size and the MIC values in ID-CAMHB were one dilution higher or lower than the suggested resistance breakpoint (2 mg/L).

The MBC/MIC ratios evaluated by the following standard criteria [16] showed a bactericidal activity for FDC in all strains but one (*A. baumannii* ATCC 19606).

### Population analysis profile (PAP)

In addition to previous determinations, PAP analysis was performed for all of the study strains in order to determine the frequency of bacterial cells growing on agar supplemented with various concentrations of FDC. Results are reported in Table 1b and Figure S1. Eight out of ten strains were heteroresistant as the number of colonies growing at antibiotic concentrations up to 16 times the breakpoint (that is, not just at 2-4 times as stated in the guidelines) were at least 1:10^6^ of those growing on antibiotic-free plates. Abau5 was considered resistant because the colonies growing in the presence of 2 mg/L of FDC outnumbered 50% of colonies growing on antibiotic-free plate, vice versa *A. baumannii* ATCC 17978 was considered fully susceptible (Figure S2). Moreover, all strains but *A. baumannii* ATCC 17978 grew on MHA plates with 32 mg/L of FDC in the form of very tiny colonies (Figure 1a), and microscopic examinations of these colonies revealed the presence of filamentous bacteria (Figure 1b). These colonies were slightly visible after 24 hours of incubation, becoming big enough to be enumerated after 48 hours. An internal control experiment revealed no differences in the colony growth on MHA plates with FDC previously incubated at 37 °C for 48 hours before inoculation and plates inoculated right after their preparation, suggesting FDC stability at 37 °C for at least 2 days. Small colonies immediately turned back to their original shape once the drug was removed (image not shown). These data confirm the caveat of usual standard laboratory testing to detect heteroresistance, very widespread in our isolates.

**Figure 1.**
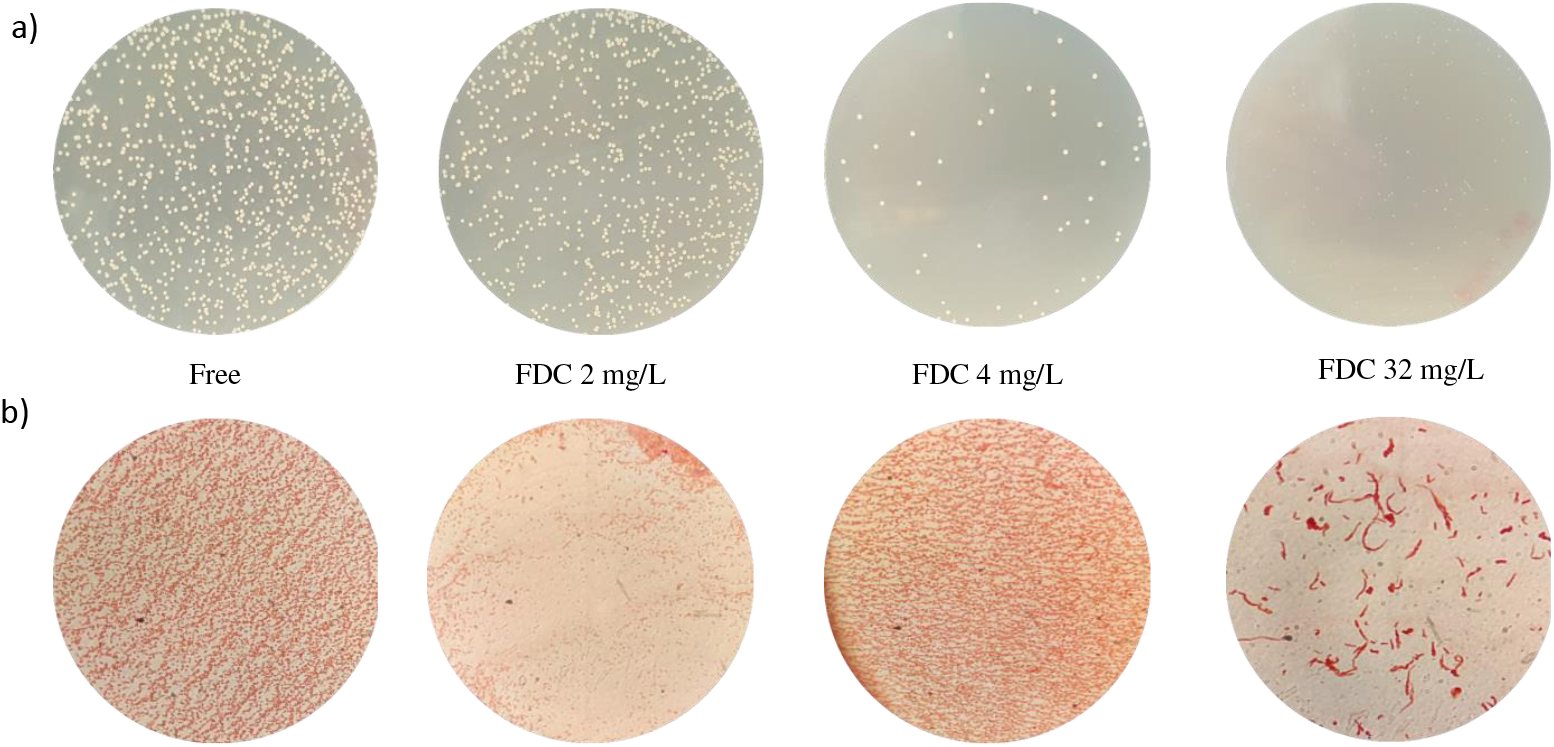
Representative model of small colonies growing and microscopic shape of non-susceptible or heteroresistant strains of *A. baumannii* at cefiderocol concentration higher than their MIC (ACICU – MIC: 2mg/L).

### Regrowth and stability of non-susceptible induced isolates

During MBC experiments, 4 strains showed a regrowth of approximately 10^3^ cells at the highest FDC concentration of 32 mg/L despite proving susceptible to FDC by DD testing and the drug demonstrating *cidal* activity (Table 2).

**Table 2.**
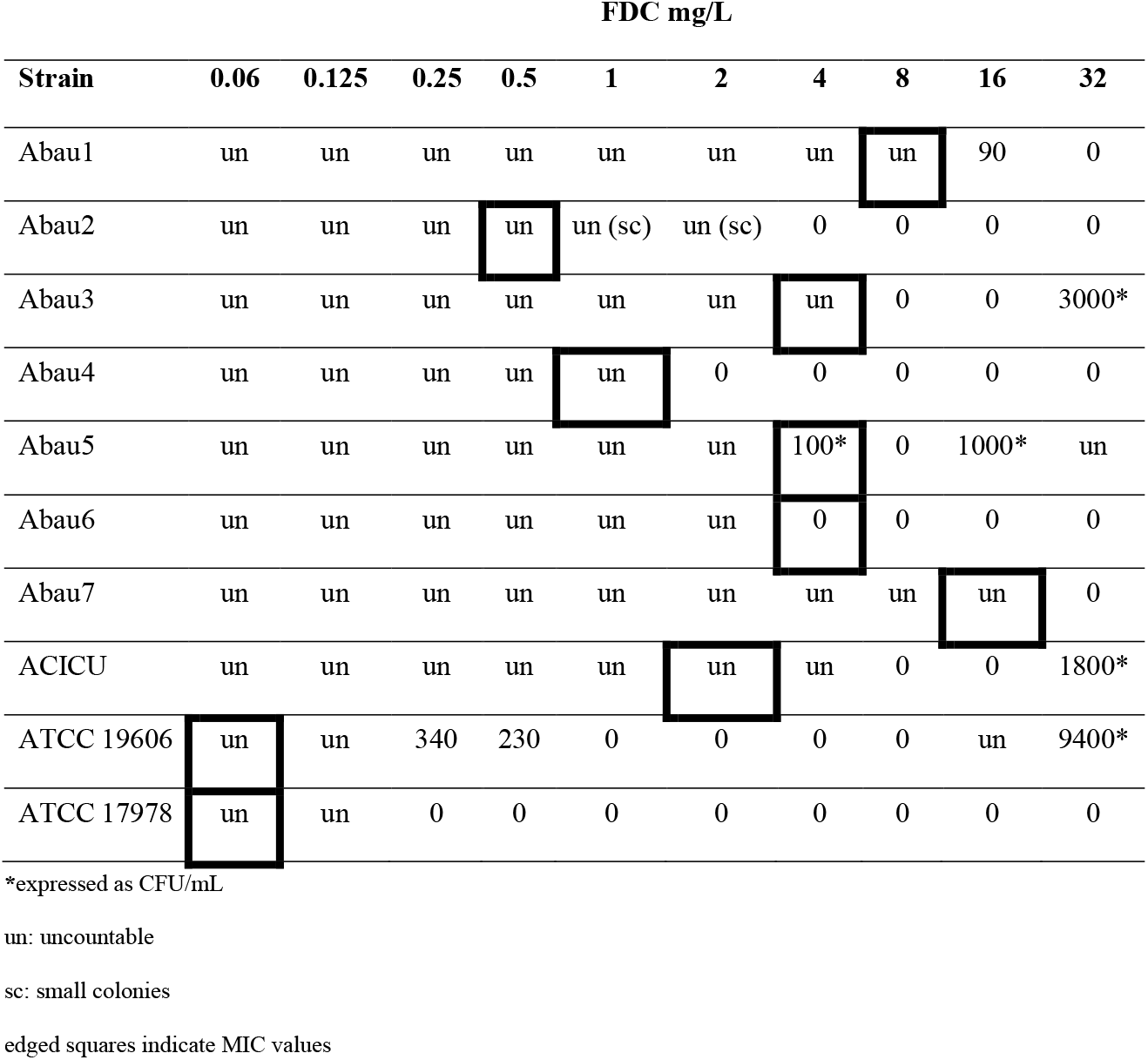
Regrowth from MBC/MIC assays.

| Strain | FDC mg/L |  |  |  |  |  |  |  |  |  |
| --- | --- | --- | --- | --- | --- | --- | --- | --- | --- | --- |
|  | 0.06 | 0.125 | 0.25 | 0.5 | 1 | 2 | 4 | 8 | 16 | 32 |
| Abau1 | un | un | un | un | un | un | un | un | 90 | 0 |
| Abau2 | un | un | un | un | un (sc) | un (sc) | 0 | 0 | 0 | 0 |
| Abau3 | un | un | un | un | un | un | un | 0 | 0 | 3000* |
| Abau4 | un | un | un | un | un | 0 | 0 | 0 | 0 | 0 |
| Abau5 | un | un | un | un | un | un | 100* | 0 | 1000* | un |
| Abau6 | un | un | un | un | un | un | 0 | 0 | 0 | 0 |
| Abau7 | un | un | un | un | un | un | un | un | un | 0 |
| ACICU | un | un | un | un | un | un | un | 0 | 0 | 1800* |
| ATCC 19606 | un | un | 340 | 230 | 0 | 0 | 0 | 0 | un | 9400* |
| ATCC 17978 | un | un | 0 | 0 | 0 | 0 | 0 | 0 | 0 | 0 |

\*expressed as CFU/mL
un: uncountable
sc: small colonies
edged squares indicate MIC values

Among the strains showing regrowth after MIC and MBC assays, Abau3 was the only one able to grow after various passages in MHA plate and in ID-CAMHB containing 32 mg/L of FDC. As expected, the FDC MIC value of the strain picked up from the MHA plate containing FDC was significantly higher than the original MIC of Abau3, i.e., MIC >32mg/L *vs* MIC 4 mg/L. Nevertheless, non-susceptibility was lost after two passages of the induced Abau3 strain in antibiotic- free plates (Table 3). A DD test on induced Abau3 gave an inhibition zone of 6 mm, confirming its non-susceptible profile, while the same test performed on Abau3 that had lost induction yielded an inhibition zone of 18 mm, as typical for susceptible strains (Table 3).

**Table 3.** Induction and maintenance of cefiderocol non-susceptibility.

| Strain | MH Agar |  | ID-CAMHB |
| --- | --- | --- | --- |
|  | DD mm | S/NS<sup>‡</sup> | MIC mg/L |
| Abau3 post-induction with 32 mg/L<br>from a 32 mg/L FDC agar plate | 6 | S | >32 |
| Abau3 post-induction with 32 mg/L<br>from a FDC-free agar plate | 18 | NS | 2 |

<sup>‡</sup>characterized as susceptible or non-susceptible in accordance with the EUCAST breakpoints for cefiderocol and AST: a zone diameter $\geq 17$ mm is typical for isolates with MIC values $\leq 2$ mg/L
(*Enterobacterales*, *P. aeruginosa*, and PK/PD breakpoint) [14]
i.e.: colonies within the zone of clearing
S: susceptible; NS: non-susceptible

Microscopic investigations of induced Abau3 showed no or very little differences in the bacterial cells grown in the presence or absence of FDC, with only a few filamentous bacteria in the former condition (Table 3).

Effect of β-lactamase inhibitors on cefiderocol susceptibility restoration

As our strains harbored multiple classes of β-lactamase resistance genes, the addition of BLIs to FDC could to some extent restore its activity. The inhibition zones of CAZ and CZA for the non- susceptible induced Abau3 strain grown on antibiotic-free MHA plates and plates with 32 mg/L of FDC are shown in Figure 2. The lack of inhibition around the disks containing CAZ and CZA on antibiotic-free MHA plates indicated that the strain was resistant to ceftazidime and that avibactam would not increase its efficacy against the resistant strain. Conversely, the emergence of an inhibition zone around the CZA disk, together with its absence around the CAZ disk, on MHA plates supplemented with FDC revealed that avibactam was able to restore FDC antimicrobial activity. SAM and AMP strips did not show any difference in plates with or without FDC, indicating a poor activity of sulbactam in restoring the efficacy of FDC in this induced isolate.

**Figure 2.**
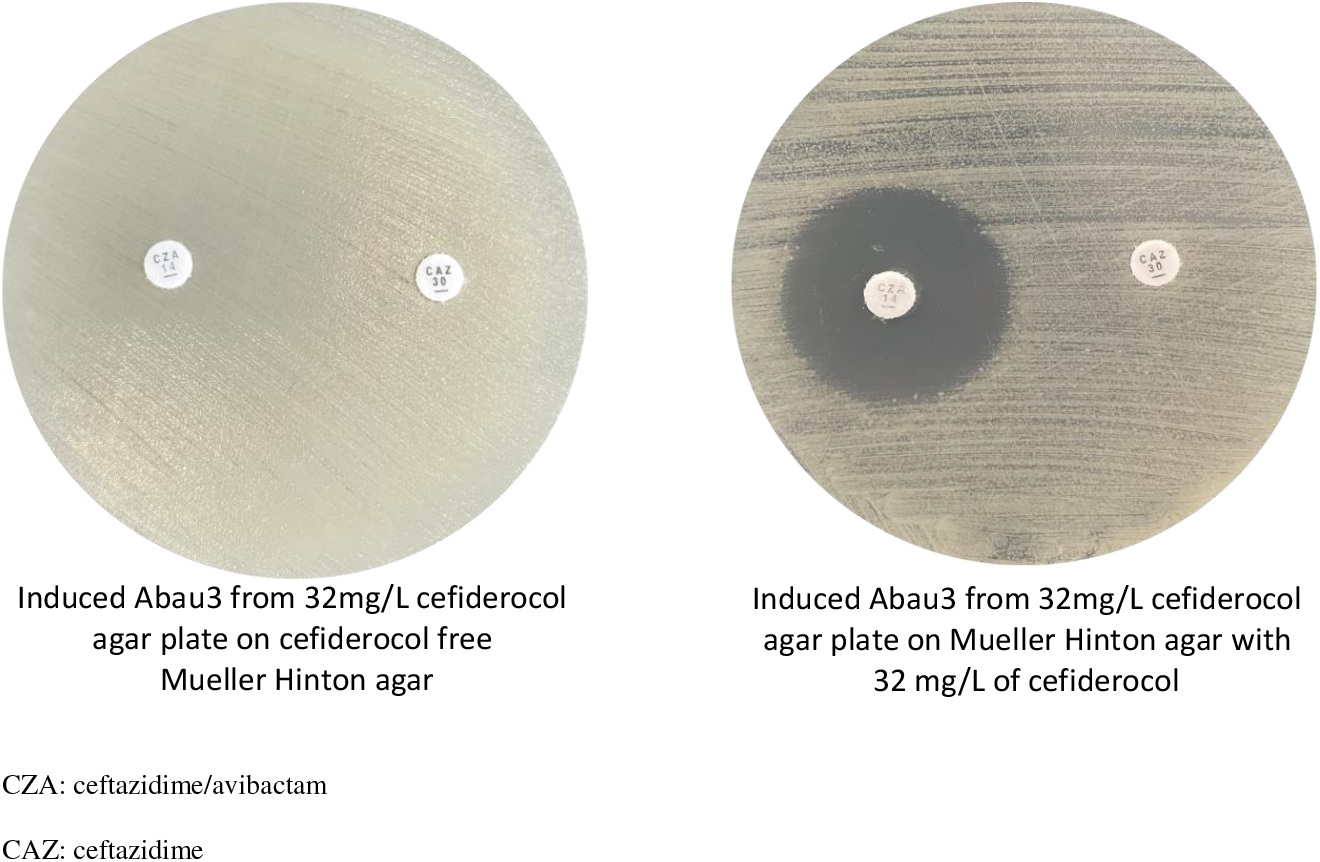
Effect of avibactam on cefiderocol susceptibility regain.

The same tests performed on the other strains with FDC MIC >1 mg/L (i.e., Abau1, Abau3, Abau5, Abau6, Abau7, and ACICU) revealed different activities for avibactam and sulbactam regardless of their FDC MIC values. Particularly, avibactam restored FDC activity for Abau3 and Abau7, while sulbactam restored FDC activity for the Abau3, Abau5, Abau6 and Abau7 strains (resistant to SAM with MIC values >256 mg/L).

## DISCUSSION

Cefiderocol, a novel siderophore cephalosporin, gained great attention due to its peculiar mechanism of entry into cells (the so-called Trojan horse-like approach [4]) and undoubted activity against MDR organisms, including *A. baumannii* [3,19,20]. In this context and in parallel with the increased expectations of its use, difficulties emerged in terms of: i) defining the best method to be used for AST, resulting from the peculiar requirement for the drug to be assessed in ID-CAMHB in order to induce siderophore-mediated entry; and ii) some cases of clinical and microbiological failures reported in the literature, related to evoked heteroresistance to this drug [5,21].

From what is nowadays known, major gaps are still to be filled in all these fields.

Our study aimed to test cefiderocol susceptibility in some strains of *A. baumannii*, both clinical and reference strains, focusing initially on the conditions for susceptibility testing by the use of two different liquid media and the disk diffusion method to determine the level of reproducibility of each set of testing; further, we aimed to determine the *cidality* of the drug, followed by PAP analysis experiments, to evaluate the presence of resistant cell subpopulations as well as the activity of BLIs in restoring cefiderocol’s antibacterial activity.

The *A. baumannii* strains included in the study belong to two different ST types and were MDR, including resistance to meropenem (9/10 of the sample) and, to a lesser extent, colistin (2/10 of the sample). Resistome analysis by NGS demonstrated a plethora of OXA and ADC variants in their genomes. In these strains, FDC showed a different degree of susceptibility when a disk diffusion breakpoint ≥17 mm was used (5 strains were susceptible and 5 non-susceptible or heteroresistant due to the presence of colonies within the inhibition zone). The geometric mean of the MIC values performed at least three times in CAMHB and ID-CAMHB correlate better with the disk diffusion results whenever the DD and MIC values were distant from their respective breakpoints (17 mm and 2 mg/L, respectively); instead, every time those values coincided or were very close to the breakpoints, the rate of discordance between the two methods was higher. The same discordance can be found in the EUCAST MIC-zone diameter correlates for *A. baumannii* for strains with a DD diameter between 16 and 18 mm (Area of Technical Uncertainty, ATU) [15].

Considering the MBC/MIC ratio, even if FDC proved bactericidal in almost all isolates, *A. baumannii* seemed intrinsically prone to adapt to increased drug concentrations, showing a general ability (four out of ten strains) to survive at the highest drug concentration.

In particular, these subpopulations exhibited smaller colonies and a change in the shape of the microorganisms, from a coccobacillary form to filamentous cells. The regrowth capacity of several tiny colonies, as demonstrated at the highest concentrations of FDC, retested immediately after isolation and after two passages in antibiotic-free medium, was found to be unstable. Indeed, all colonies reverted to the original MIC value. The change in the shape of the microorganisms could be explained by the mechanism of inhibition of FDC, which binds PBP3 as the most important target of the drug, followed by PBP2 and 1 [22], as well as by the propensity of *A. baumannii* to change its colony morphology and cell shape in response to many stress factors [23]. However, the different resistance profiles in our strains seem to be not related to PBP3 integrity as there were no amino acid changes in 7 samples when compared to the fully susceptible ATCC 17978 strain and only one or two non-synonymous mutations in the remaining 2 isolates. To note, the H370Y amino acid change found in ACICU have been previously reported by Nordmann et al. arising in an *A. baumannii* strain following the treatment with FDC that resulted in the increase of the MIC value from 1 to 4 mg/L [24].

Regrowth at the highest concentration and its loss after as little as two passages without FDC confirm the adaptability of this species to the surrounding environment, responding to a myriad of intra- and extra-cellular signals [25]. All these regulatory circuitries, widely represented in the genome of this microorganism, were evoked to explain the emergence of resistant cells expressing distinct and critical phenotypes - for example, becoming resistant - in order to survive and adapt to hostile situations.

The same phenomenon of regrowth at increased FDC concentrations in populations greater than 1×10^6^ CFU/ml was observed during population analysis profiling (PAP). Of the 10 strains tested, only one was frankly susceptible and one frankly resistant; all others could be defined as heteroresistant to the drug. Even in this case, tiny colonies and morphological changes were observed. In our experiments, very tiny colonies started to grow right after the first overnight incubation but were too small to be counted, and they continued to grow larger up to 48 hours of incubation to reach a size which allowed us to count and pick them up. Once again, our *A. baumannii’s* FDC susceptibility profile was inconsistent when analyzing PAP results. In fact, only the frankly resistant and the frankly susceptible strains were classified in the same way based on DD and MIC evaluations. Despite being quick and easy to perform, these latter methods are not suited to determine the presence of unstable heteroresistant subpopulations that could be the reason for some therapeutic failures with FDC and other antibiotics. These cases need to be further investigated by PAP to better classify *A. baumannii* resistance profile, as suggested by Band et al. for *Enterobacterales* [17].

Luckily, the combination of FDC with a β-lactamase inhibitor – such as avibactam or sulbactam - seems to restore FDC antibacterial activity even at concentrations several times lower than its MIC, as already shown by other authors [26]. It remains unclear why, in some strains, the synergistic activity is provided by both the tested BLIs while, in other strains, only one inhibitor restored cefiderocol’s activity. Furthermore, no correlation is evident between FDC MIC values and the activity of BLIs or the presence of different *bla*OXA genes.

## CONCLUSIONS

The need for new antimicrobial agents to address the treat of antibiotic resistance is recognized worldwide and, in this scenario, new molecules as cefiderocol are a breath of fresh air. Notwithstanding, any new antibiotics need to be thoroughly investigated in order to characterize their strengths and weakness and, perhaps more importantly, to unveil antibiotic/bacterial species interconnections.

Cefiderocol demonstrated promising activity against Gram-negative bacteria, including those referred to as ESKAPE, for which the therapeutic options are limited, such as *A. baumannii*. By investigating the resistance profile of some clinical and laboratory-adapted *A. baumannii* strains to cefiderocol, we found that these strains are not easily classified into the commonly used categories of susceptibility - especially strains with susceptibility levels very close to the proposed resistance breakpoints – due to a high prevalence of heteroresistant subpopulations. Heteroresistance in *A. baumannii* seems to be common as a stress response mechanism, but - luckily - it is apparently an unstable and transient trait. Our results, in accordance with recently published clinical observations [12,27], go in the direction of combination therapy when FDC is used to treat severe *A. bauamanni* infections.

## Acknowledgments

We would like to thank Shionogi & Co., Ltd for supplying us Cefiderocol powder, disks and strips. The authors wish to thank Prof. Paolo Visca for providing ACICU strain. We also wish to thank PharmaTranslated (http://www.pharmatranslated.com/) and in particular to Silvia Montanari for the language revision.

## Author contribution

Stefania Stefani (SS) and Stefano Stracquadanio (SStr) conceptualized the work and wrote the manuscript; Andrea Mar provided clinical isolates; SStr, CB AM and Alessia Mirabile (AMir) performed the experiments; DB, GFP, DAB and PGB made genome sequencing and mutational analysis; SS, SStr and CB analyzed the data, SS supervised and acquired the funds. All authors have read and agreed to the published version of the manuscript.

